# Young Children Integrate Current Observations, Priors and Agent Information to Build Predictive Models of Others’ Actions

**DOI:** 10.1101/365361

**Authors:** E. Kayhan, L. Heil, J. Kwisthout, I. van Rooij, S. Hunnius, H. Bekkering

**Keywords:** predictive models, development, pupil dilation, prediction error, causal Bayesian networks

## Abstract

From early on in life, children are able to use information from their environment to form predictions about events. For instance, they can use statistical information about a population to predict the sample drawn from that population and infer an agent’s preferences from systematic violations of random sampling. We investigated how young children build and update models of an agent’s sampling actions over time, and whether a computational model based on the causal Bayesian network formalization of predictive processing can explain this process.

We formalized three hypotheses about how different explanatory variables (i.e., prior probabilities, current observations, and agent characteristics) are used to build predictive models of others’ actions. We measured pupillary responses as a behavioral marker of ‘prediction errors’ (i.e., the perceived mismatch between what one’s model of an agent predicts and what the agent actually does), as described in the predictive processing framework. Pupillary responses of 24-month-olds, but not 18-month-olds, showed that young children integrated information about current observations, priors and agents to generate predictive models of agents and their actions.

These findings shed light on the mechanisms behind toddlers’ inferences about agent-caused events. To our knowledge, this is the first study in which young children’s pupillary responses are used as markers of prediction errors, and explained by a computational model based on the causal Bayesian network formalization of predictive processing. We argue that the predictive processing framework provides a promising explanation of the way in which young children process other persons’ actions.

**Highlights:**

- We present three formalized hypotheses on how young children generate predictive models of others’ sampling actions.
- We measured pupillary responses of children as a behavioral marker of prediction errors as described in the predictive processing framework.
- Results showed that young children integrated information about current observations, prior probabilities and agents to generate predictive models about others’ actions.
- A computational model based on the causal Bayesian network formalization of predictive processing can explain this process.

## Introduction

From a very young age, children are able to infer that some events are more probable than others. They use these inferences to form expectations about future events and show surprise when these events unfold differently. When 12-month-old infants see a container in which three identical objects and a single object move around before one of them exits the container, they look longer when the single object leaves the container rather than one of the majority objects (Téglás, Girotto, Gonzalez, & Bonatti, 2007). A similar situation occurs when young children observe more or less probable actions of another person. Xu and Garcia (2008) showed that infants as young as eight months of age look longer at a sample of colored balls if it is picked from a population of balls with mostly other colors, suggesting that they were expecting a different sample given the population. In other words, if a population contains mainly red balls, infants expect sampling from the population to be random, resulting in a sample of mainly red balls. If this expectation is violated, this is reflected in an increased looking time. If, however, an agent consistently picks the same items from a population in a non-random way (e.g., all white balls from a predominantly red population), the observer might interpret this as an indication of a preference for a certain item. In a study by Kushnir, Xu, and Wellman (2010), 20-month-old children observed an agent picking five toys of the same type from a population box that held mostly toys from a different type. When the toddlers were then asked to give the agent the toy he liked best, they often chose the toy that the agent picked before. If, on the other hand, the agent had picked the same toys from a population box in which the two types of toys were more evenly distributed, they picked this toy less often. These results suggest that infants infer preferences of others based on violation of random sampling.

Previous research has also shown that infants use probabilistic information to inform their predictions about others’ actions. For example, Henrichs and colleagues (2014) investigated whether goal certainty modulated action prediction patterns of 12-month-old infants. In an eye-tracking paradigm, infants observed hands reaching towards one of three objects on the table, grasping the objects and placing them in a bowl. Infants performed earlier gaze shifts in the frequent condition when the hand reached for the same object in all trials as compared to the non-frequent condition, in which the hand reached for different objects across trials. These findings indicate that infants use probabilistic data to make predictions about others’ actions.

The idea that inferences about the probability of events are used to generate predictions about these events is in line with the predictive processing framework. According to this framework, we build generative models to predict incoming information (Clark, 2013; Friston, 2005). This predicted input is compared to the actual input, and the difference (i.e., the prediction error) is used to update the predictive models. Previous studies have showed neural markers of prediction errors (e.g., Egner, Monti, & Summerfield, 2010; den Ouden, Daunizeau, Roiser, Friston, & Stephan, 2010; Phillips, Blenkmann, Hughes, Bekinschtein, & Rowe, 2015; van Pelt et al., 2016; Wacongne et al., 2011). Other studies suggest that prediction errors can also be assessed through measurements of pupillary responses, as these have been shown to correlate with prediction errors in a predictive-inference task (Nassar et al., 2012) and with reward prediction errors (Preuschoff, ’t Hart, & Einhäuser, 2011) in adults.

Because pupillary responses occur involuntarily without explicit instructions, the method has also been valuable to unravel perceptual and cognitive processes in preverbal infants (Hepach & Westerman, 2016; Laeng, Sirois & Gredebäck, 2012). Particularly, pupil dilation has been assumed to represent infants’ violations of expectations (Gredebäck & Melinder, 2011; Jackson & Sirois, 2009) For example, Addyman, Rocha, and Mareschal (2014) investigated time perception in infants by using a paradigm in which recurring targets were omitted. Results of this study showed that 4- to 14–month-old infants showed increased pupil dilation to the absence of the targets at anticipated time intervals indicating a violation of their expectations of interval timing. In the present study, we take this approach one-step further and use infants’ pupillary responses as indirect behavioral markers of prediction errors in young children.

If young children build a predictive model of other people’s actions in which the probability of events is represented, then we would expect to see increased pupil dilation when children observe improbable events. Looking times may be expected to increase in a similar way. Although they are indeed the most widely used measure of prediction violations (Aslin, 2007), they are often measured over relatively long periods after stimulus presentation (e.g., > 12 sec in Wellman et al. (2016); > 5 sec in Xu & Garcia, 2008; > 6 sec in Xu & Denison, 2009). As such, it is difficult to distinguish between initial time-locked responses to the violation of predictions and cumulative responses that might reflect post-hoc processes (Jackson & Sirois, 2009). The shorter time scale at which pupillary responses occur may provide unique insights into how different explanatory variables are used to form predictions over time. In order to explain the mechanism behind infants’ responses, we compared the qualitative pattern of pupillary responses to prediction error patterns as generated by a predictive processing model.

In the current study, we investigated how 18-month-old infants and 24-month-old toddlers build predictive models of others’ actions. Do they, as suggested by previous studies, integrate information about prior probabilities and current observations to predict a subsequent event? Moreover, is the fact that the agent performing the action might have a bias also taken into account? In an experiment in which young children observed an agent performing a series of more or less probable sampling actions, we analyzed the changes in pupillary responses over trials to examine in a fine-grained manner how infants and toddlers build a model about an agent’s sampling actions. Here it is important to note that differently from previous studies (e.g. Xu & Garcia, 2008; Xu & Denison; 2009) we measured the pupillary responses after each sampled outcome separately. This unique feature of our design allowed us to monitor how infants’ predictions and the resulting prediction errors evolved over time.

We created an experimental setting in which children observed a puppet drawing balls from a population box and placing them in a row in an open container one by one. The population box contained balls of two colors, with a ratio of 1:4. In the *Minority-first* condition, the puppet first performed a series of improbable actions by picking four colored balls of the minority color, before picking a ball in the majority color. In the *Majority-first* condition, the puppet performed a series of more probable actions by drawing four balls of the majority color, followed by one minority color ball.

Our computational approach allowed us to generate distinct hypotheses prior to data collection. We formalized three hypotheses to explain how young children build predictive models of others’ sampling actions. These formalized hypotheses (of which the precise computational characterization is presented in the Appendix) provided us with a predicted pattern of prediction errors in all trials and conditions. We assume that in our experimental setup, there are three variables that are used to model the environment: (1) the previous observations (i.e. the balls that were sampled before), (2) the prior probabilities of an event (i.e. the probability of balls of a certain color being picked given the relative number of balls in a population), (3) the agent’s biases (i.e. tendency to pick a certain color). These variables increase in terms of level of complexity: whereas the first one is dependent on change detection, the second one relies on statistical inference, and the third one is based on processing of unobservable agent information. The hypotheses differ with respect to whether and how these variables are taken into account.

According to the first hypothesis, children build a predictive model solely based on the sample drawn so far. In other words, if a green ball were drawn from the population before, the best prediction of the next event would be that another green ball would be drawn. As evidence builds up, this prediction gets stronger, resulting in decreasing prediction errors over the course of trials. However, when the last ball differs from the previous ones, this should lead to an increase in prediction errors. As a result, over time, one would observe a decrease in pupillary responses in both the *Minority-first* and *Majority-first* conditions, as several balls of the same color are being picked repeatedly (see Figure 1A). However, when the last ball differs from the previous ones, one would expect the pupillary response to increase for the last sampling event as compared to the previous event in both conditions. In the first hypothesis, information about the fact that the ball was sampled from a population with a specific distribution or sampled by an agent who may have certain biases is assumed not to be taken into account.

**Figure 1:**
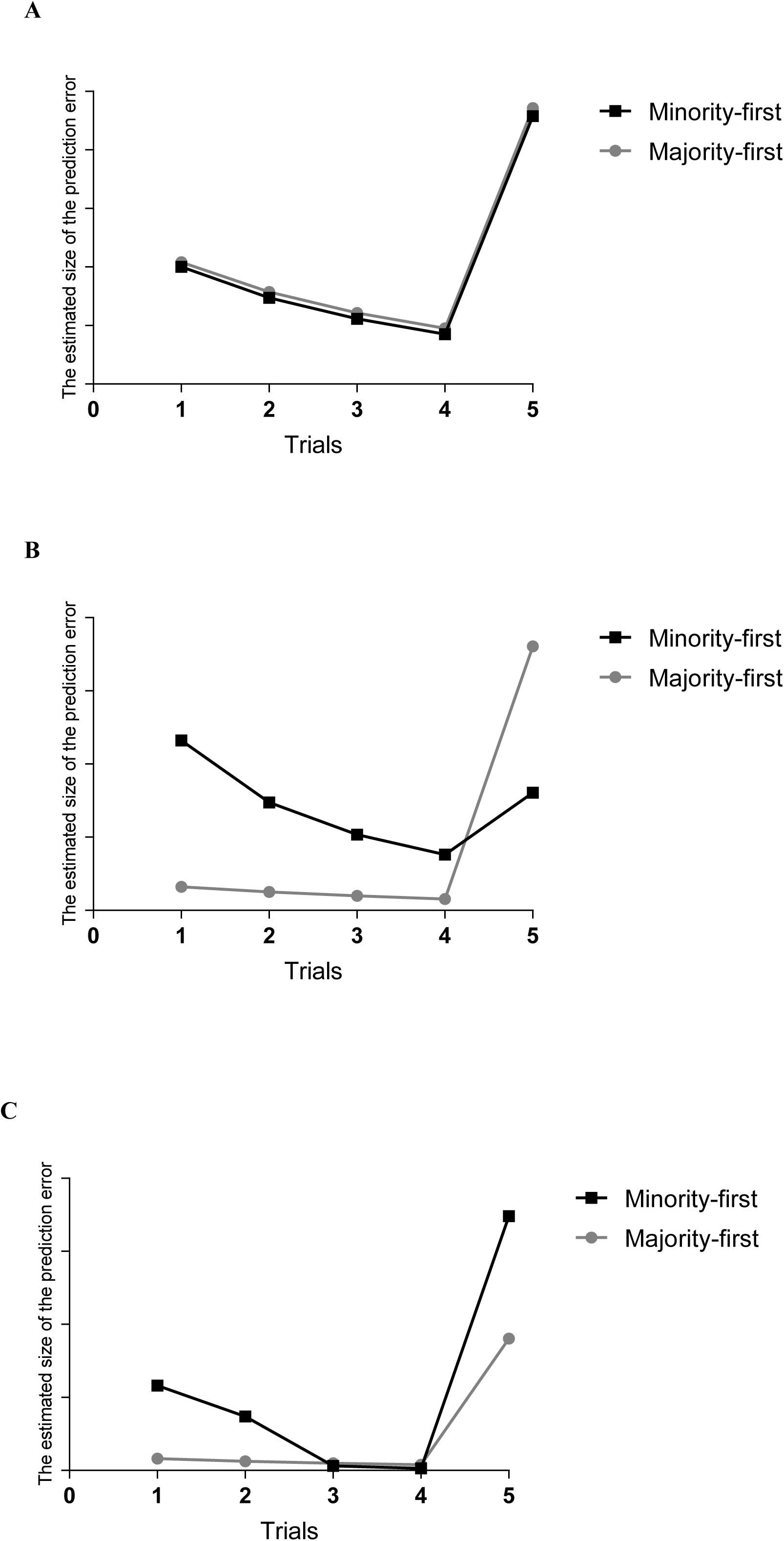
The estimated size of the prediction error (and pupil dilation thereof) for the two experimental conditions as predicted by the computational models, based on Hypothesis 1 (A), Hypothesis 2 (B), and Hypothesis 3 (C).

The second hypothesis predicts that children do take probabilistic information about the population into account, but still ignore the information about the agent. This hypothesis entails that children use both prior probability and current observations to predict the next sampling event: a green ball picked from a mostly yellow population is improbable, but becomes slightly more probable after it happened a few times. Based on this hypothesis, one would expect the pupillary response for the first two trials to be large for the *Minority-first* condition. However, when balls of the same color are picked repeatedly, the pupillary responses should decrease in the subsequent trials. Yet, when the last ball differs from the previous observations but it is more probable given the distribution of colors in the population box, we expected the pupillary responses to increase only slightly from the fourth to the fifth sampling event. In the *Majority-first* condition, on the other hand, one would expect the pupillary responses to be lower in the initial trials, as compared to the *Minority-first* condition, given that it is more probable to pick yellow balls from the given population box. However, because the last ball is different from the previous ones and it is less probable given the ratio of balls in the population box, we predicted a larger increase in the pupillary responses from the fourth to the fifth sampling event in *Majority-first* condition as compared to the *Minority-first* condition (see Figure 1B).

According to the third hypothesis, children do not only process the sampling actions as probabilistic events, but they also consider unobservable variables such as an agent’s bias while doing so. They integrate the prior probability and current observations in a way that includes agent characteristics as an explanatory variable that predicts observed actions. If an agent consistently performs an improbable action, then sampling might not be random but driven by some characteristics of the agent (e.g., a bias for picking a certain color). In terms of experimental findings, this hypothesis predicts that children show larger pupillary responses in the *Minority-first* condition, as compared to the *Majority-first* condition, during the first trials, because it is less probable to pick the minority balls repeatedly from the population box. Then, as several minority balls are selected in a row in the *Minority-first* condition, the joint probability of the events as a whole becomes so low that children will update their models. As a result, pupil dilation should decrease after the first few trials, as they will then assume that the agent deliberately selects balls in minority colors because of a picking bias and will expect the agent to keep doing this, consistent with this picking bias. However, as the last ball differs from the first four, their predictions based on the updated model will be violated and there will be a large increase in the pupillary responses again in this condition. On the other hand, in the *Majority-first* condition, there is no reason to reject the assumption that the agent samples randomly. Therefore, neither the fact that majority colors are being picked in a row nor the color of the last ball deviates from the previous ones is too surprising: the distribution of colors in the sample is consistent with the distribution in the population. If this is the case, the pupillary responses in the first few trials in *Majority-first* condition will be lower than in the *Minority-first* condition and they will only slightly increase in the last trial as compared to the previous trial, as the last balls differs from the previous observations (see Figure 1C).

Children’s predictive models of other’s actions may become more precise over the course of development. For example, they might get more precise in representing statistical information, as they get older. It could also be that given the increased amount of social experience, they might become more proficient in recognizing others’ preferences. Indeed, developmental research on social cognition suggests that children’s attributions in social situations change as they accumulate more statistical evidence about agents in different situations through experience (Meltzoff & Gopnik, 2013; Gopnik & Wellman, 2012). For example, Ma and Xu (2011) investigated whether 24-month-old toddlers and 16-month-old infants use statistical information to infer that others might have preferences different from their own. Although 24-month-olds first assumed that the experimenter would share their preference for a certain object, they were able to revise this assumption when the experimenter repeatedly chose another object, only if sampling appeared to be non-random. Whereas 24-month-old children were able to infer that the experimenter had a preference different from their own, 16-month-old infants showed weaker evidence for such an inference. As Ma and Xu (2011) argue, these findings suggest that the ability to reason about the subjective nature of preferences develops between 16 months and 2 years of age. Given the previous literature, we included two age groups (24- and 18-month-olds) in the current experiment to investigate if the use of probabilistic and agent information in generating predictive models changes between 18 and 24 months.

The hypotheses described earlier were formalized based on the Predictive processing framework. Their computational elaboration can be found in the Appendix. These formalizations provide us with an estimation of prediction errors in different trials and conditions. We compared these predicted patterns of results to the actual changes in pupil dilation for both age groups. With this, we provide more insight into the way in which children use prior probabilities, current observations and agent information to build predictive models of others’ actions, and how they revise their models over time.

## Method

### Participants

We tested 48 18-month-old infants (*M* = 18 months 2 days; range 17 months 11 days–18 months 13 days) and 57 24-month-infants (*M* = 24 months 1 day; range 23 months 4 days–24 months 15 days) for the study. Ten 18-month-olds and six 24-month-olds did not complete the testing session due to fussiness. Participants were recruited from a database of volunteer families. The local Social Science Faculty’s ethical committee approved the study. All children were born full-term and had no reported developmental delays. Participating families received a book or 10 Euros in return.

### Stimuli

We created familiarization and test movies that showed animal hand puppets sampling colored balls from a population box and placing them one by one in an open container. The population box was transparent in front and had a white opaque cover on top in order to hide the sampling action. This part also served as an occluder to keep the puppets out of sight when they left the scene each time after they had drawn a ball from the population box. An opaque tube that was attached to the box led to the container on the left side of the box (see Figure 2 A-C).

**Figure 2:**
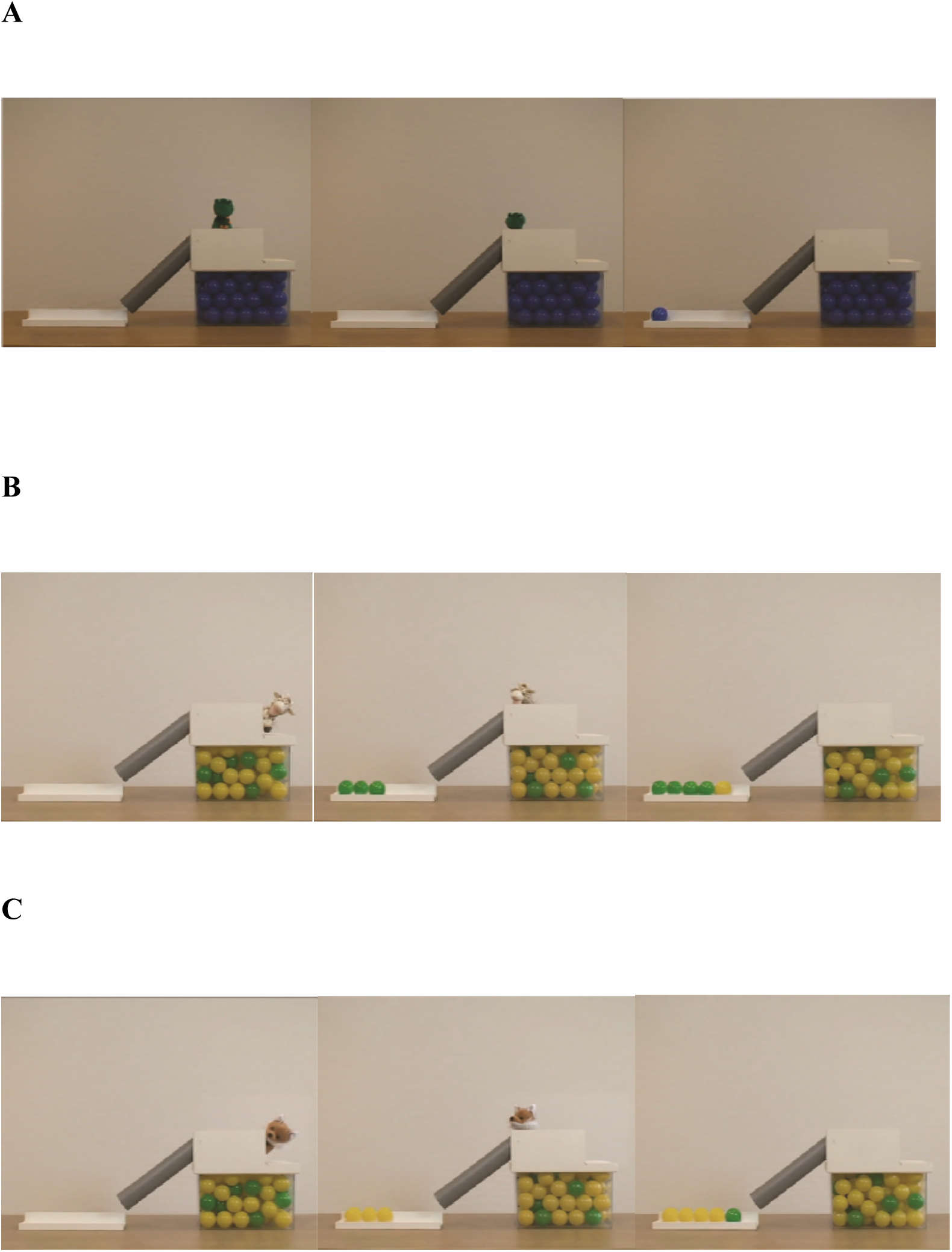
Snapshots from stimulus movies of the (A) Familiarization, (B) *Minority-first* condition, and (C) *Majority-first* condition

#### Familiarization movie

In the familiarization movie (Figure 2A), a frog puppet popped up from behind the occluded part of the population box and started an introductory talk dubbed by a female voice. The puppet introduced itself and presented the population box, the tube and the container to the child. It then popped down to pick a ball from a population box filled with only blue balls before appearing again and moving towards the tube. The puppet’s hands and the ball were hidden behind the opaque part on top of the box until it put the ball into the tube. The puppet then quickly left the scene so that distraction was minimized during the measurement of the pupillary response in test trials. Immediately after the puppet disappeared, a rolling sound was played and the ball appeared on the left side of the container. The rolling ball was not shown to the viewer in order to ensure that participants did not see its color until it appeared in the container. The puppet then popped up again and explained that the balls roll all the way down to the end, in order to familiarize the child with the sudden appearance of the balls in the container. This process was repeated for five times and took 2.07 minutes in total.

#### Test movies

Four different animal hand puppets, dubbed by two male and two female voices, were used to depict the different sampling events. We filmed each puppet for the *Majority-first* as well as for the *Minority-first* condition to counterbalance the sampling events and the associated agents across participants. Test movies (see Figure 2 B-C) were similar to the familiarization movie in terms of the set-up. However, the population box was now filled with balls in two different colors. In all movies, there was a 1:4 ratio of green and yellow balls.

As in the familiarization movie, the puppet gave a short introduction in which it told the participants its name, explored the set-up and explained that it would now pick a ball. Then, it popped down behind the white opaque part to pick a ball. At this point, the agent was not visible but a rumbling sound was presented while the balls moved inside the population box in order to indicate that the agent was picking a ball. Because the puppet was not visible during this sampling event, it was not obvious from the way in which the sampling action was performed whether it was random or not. The puppet then popped up from behind the opaque part, carried the ball, and put the ball into the tube. During this period, the ball that the puppet picked was still not visible to the participants. Immediately after the puppet left the scene, a rolling sound started lasting for 1400 milliseconds until the ball appeared on the left side of the container. The display of the sampled ball was shown for 4000 milliseconds. The puppet repeated the same process five times, picking balls one by one from the box. In the *Minority-first* condition, children observed the puppet drawing the minority color balls from the population box four times in a row before picking one majority color ball (see Figure 2B). In the *Majority-first* condition, the puppet drew four majority color balls followed by a minority color ball (see Figure 2C). We measured changes in pupil dilation after each sampling event. *Majority-first* and *Minority-first* conditions were shown twice to all participants. However, as most participants got distracted quickly after observing the first entire sampling event, only data for this first sampling event were included in the analyses. In this way, we also ensured that carry-over effects that might occur due to the repetition of sampling sequences would not influence our data. One test movie lasted for 1.48 minutes and the entire stimulus presentation lasted for 8 minutes.

The stimulus material was edited in post-production using Final Cut Studio 3 (Apple Inc.). The movies were further edited using open source video editor software Kdenlive (version 0.9.6) to match the timing and the durations of each movie. As pupillary responses are sensitive to light effects, we paid utmost attention to luminance factors. In order to ensure that the movies had similar luminance values, we color-corrected the movies using Color software (version 1.5, Apple Inc.). The audio material for the movies was recorded and edited to match the pitch and speed of the audio material between movies using Audacity software (version 2.0.5).

### Experimental set-up and Procedure

The testing procedure was identical for both age groups. Eye movements were recorded with a corneal reflection eye-tracker (Tobii 120, Tobii Technology, Danderyd, Sweden) recording gaze data at 60 Hz using a 9-point calibration procedure. The procedure was repeated if seven or fewer calibration points were detected until data for at least eight calibration points was acquired.

To control for the external luminance effects, the natural lighting was entirely blocked and the room lights were on during the calibration and testing. In order to make sure that the environmental luminance was at a certain range during the measurements, we recorded the environmental luminance during each testing using a custom-made device attached to the eye-tracker (Atlas Scientific ENV-RGB Color Detector Probe, version 1.6, combined with Arduino hardware), and the luminance values were extracted via Arduino software (version 1.0.5). The external luminance values were between 187-195 lux across testings. Participants were seated on their parent’s lap. All participants viewed the testing material at approximately 60 cm distance.

### Measures

We measured the pupillary responses for each sampling event during the first 2000 ms after the sampled ball was visible in the container. Pupil data were analyzed using custom-made MATLAB scripts (MathWorks, Friedrichsdorf, Germany). The data were cleaned via several preprocessing steps. First, we screened for missing data points. If pupil dilation values were available for both eyes, then these were averaged in order to obtain one value per sample. In case of a missing value for one of the eyes, only the value from the other eye was used for the analyses. If the difference between the left and the right eye was larger than 1 (which is considered an indication of anisocoria: a condition characterized by unequal pupil sizes) or if average values were higher than two standard deviations from the mean, the data point was considered unreliable and registered as missing. Missing data points due to blinks were corrected using a linear interpolation algorithm, in which the maximum sample gap was set to 5. Following interpolation, the data were smoothed using median and moving average filtering in order to reduce the noise in the signal. If there were more than 60 missing samples in one trial of 2000 ms, the entire trial was excluded from the analyses. Pupil diameter changes were obtained by subtracting the average pupil diameter for each sampling trial from a fixed baseline period defined as the first 1000 ms from the start of each movie before the puppet appeared for the first time. Finally, in case a value for one trial deviated more than two standard deviations from the overall trial average, it was considered an outlier and removed from further analyses. Before computing further statistical analyses on the data, we first conducted one sample t-tests against zero for each age group to ensure that there were indeed significant changes in pupillary responses as compared to the baseline level (cf. Gredebäck & Melinder, 2011). These analyses were informative as they provided a validation check for further statistical analyses testing the effects of task manipulations on pupillary responses.

## Results

Measuring the pupillary responses after each sampled outcome separately allowed us to monitor how the children’s predictive models evolved over time. One sample t-tests for 18-month-olds showed that overall, there was no significant change in pupillary responses as compared to the baseline level (t (34) = 1.11, *p* = .27). On the other hand, 24-month-olds showed significant increases in their pupillary responses as compared to baseline level (*t* (50) = 4.18, *p* < .01). Pupillary responses of the 24-month-olds and the 18-month-olds are illustrated in Figure 3 and Figure 4, respectively.

**Figure 3:**
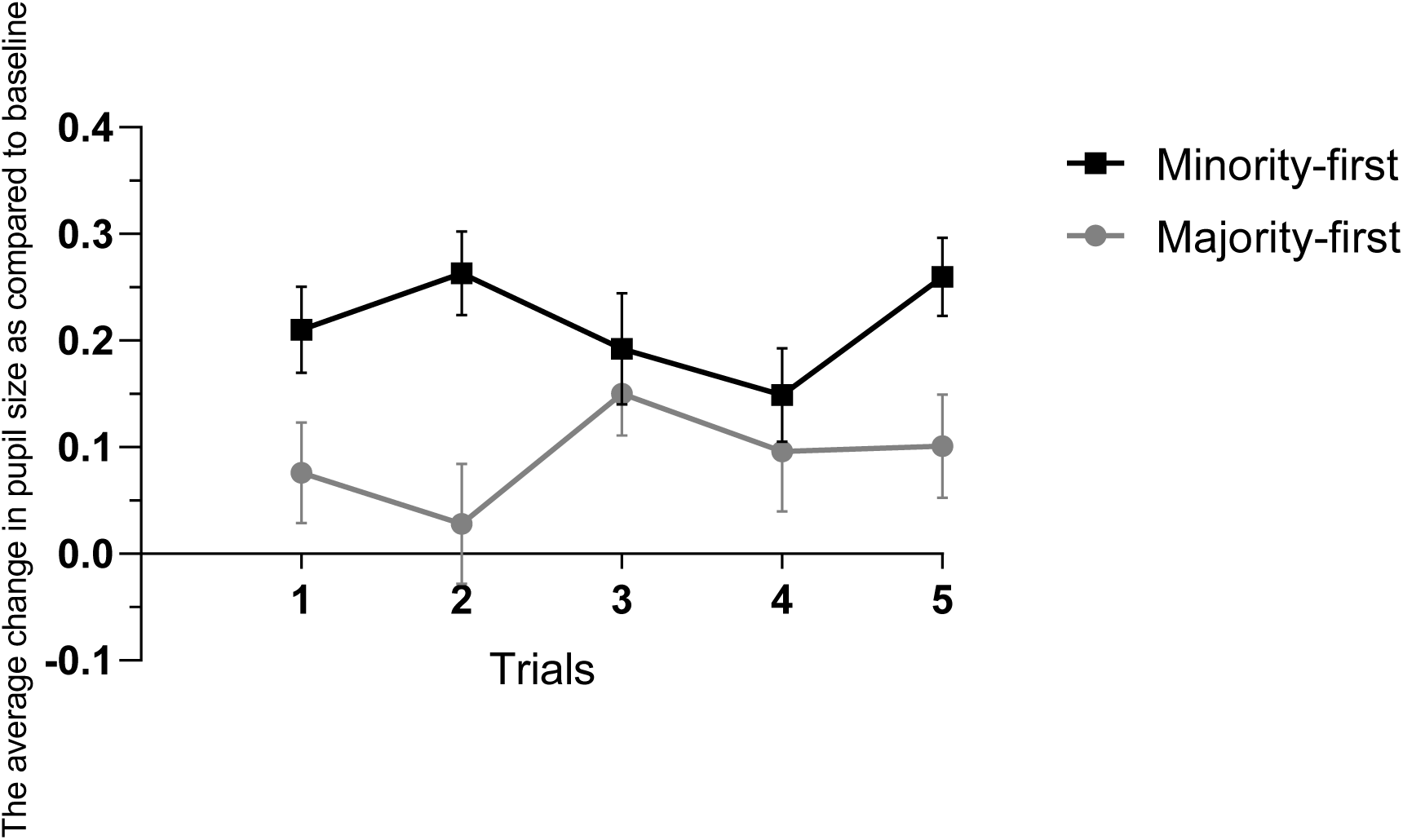
Average change in pupil size as compared to a fixed baseline period in *Minority-first* (black line) and *Majority-first* (gray line) conditions over the course of trials in 24-month-olds. Error bars represent SEMs.

**Figure 4:**
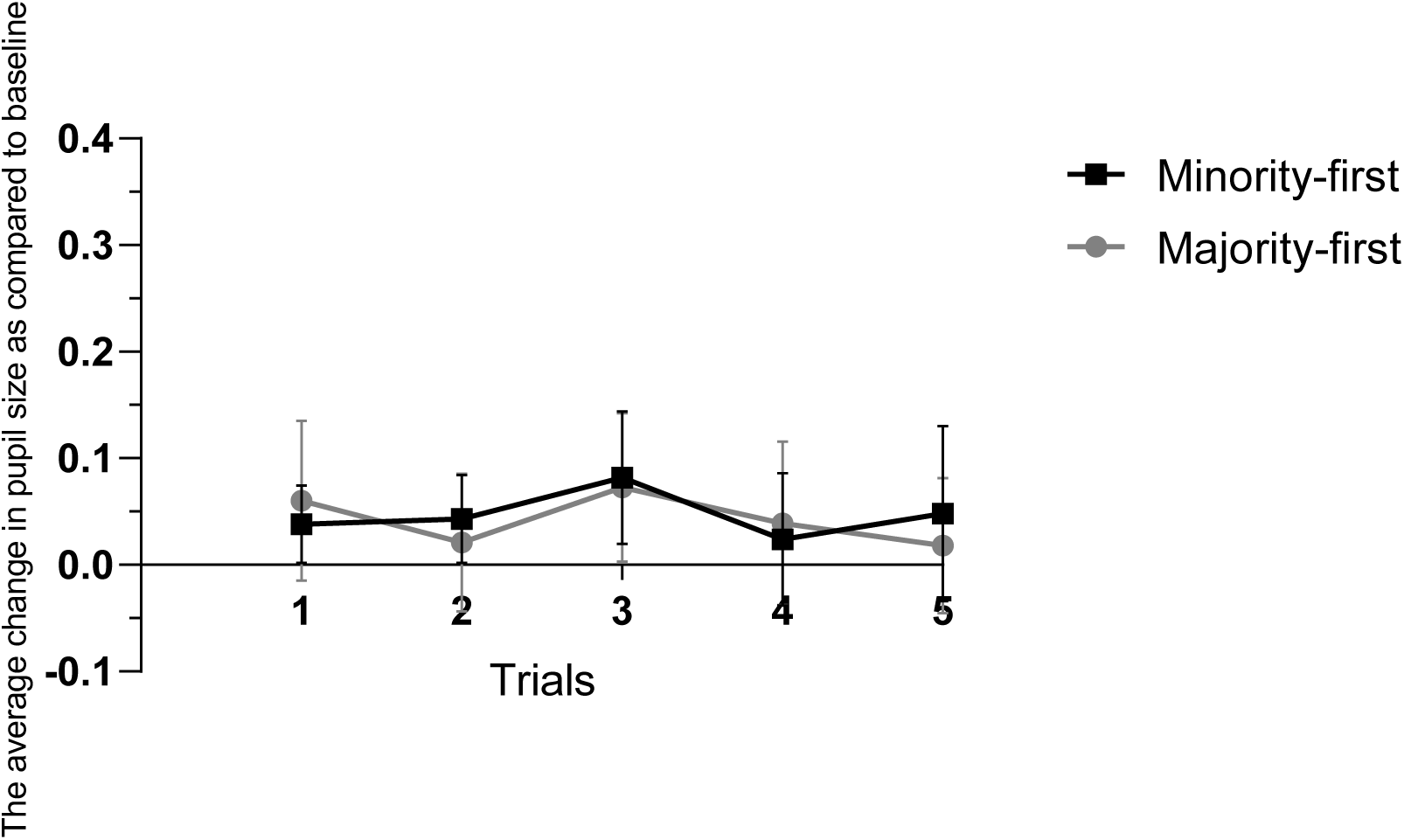
Average change in pupil size as compared to a fixed baseline period in *Minority-first* (black line) and *Majority-first* (gray line) conditions over the course of trials in 18-month-olds. Error bars represent SEMs.

Because there was no difference in pupil response as compared to the baseline level in 18-month-olds, we did not compute further analyses for this age group. We reasoned that further analyses with this age group would be uninformative if not misleading given that there were no changes in pupillary responses as compared to the baseline level, which nullify interpretations of the changes in pupil dilation as a function of different trials and conditions. As we did find a significant change in overall pupillary responses as compared to the baseline level in 24-month-olds, we focused our further analyses on this age group.

As we predicted differential response patterns for different combinations of trials given the differences in sampled outcomes, we examined the first two and last two trials separately. To examine participants’ initial responses to the probability of the outcomes, we first ran a repeated measures ANOVA with condition (*Minority-first* vs. *Majority-first)* as a between-subjects factor and first two trials (1 and 2) as a within-subjects factor. This analysis revealed a significant trial by condition interaction (F (1, 42) = 4.80, *p* = .03, *η*^2^ = 0.10). As shown in Figure 3, this interaction was mostly driven by the larger difference in pupil dilation between conditions in the second trial. Still, follow-up t-tests showed that in the first trial, pupillary responses significantly differed between the *Minority-first* condition (*M* = .22, *SD* = .20) and the *Majority-first* condition (*M* = .08, *SD* = .22), *t* (46) = 2.15, *p* = .04. Similarly, in the second trial, the difference in pupillary responses between the *Minority-first* condition (*M* = .25, *SD* = .18) and the *Majority-first* condition (*M* = .01, *SD* = .29) was significant, *t* (40.61) = 3.47, *p* < .01. These results show that in both trials, 24-month-old toddlers’ pupil dilation was larger when they observed an improbable sampling action as compared to a probable action. This finding supports the sub hypothesis of both the second and the third hypothesis: toddlers initially expected the samples to represent the distribution in the population box, thereby assuming that the agent samples from the box randomly. Moreover, as the joint probability of sampled outcomes became less probable with each sample in the *Minority-first* condition, the difference between the conditions became larger in the second trial.

When toddlers observed the agent consistently performing an improbable action (i.e., selecting minority color balls repeatedly), our third hypothesis would assume that this led them to revise their predictions: sampling might not be random, but biased towards a certain color because of some characteristics of the agent (e.g., the agent deliberately selects a certain color). After such a revision of the predictive model, children would start predicting the agent to pick the minority color ball (which was previously considered improbable) and now would be surprised to see the majority color ball appear. On the other hand, when toddlers observed the agent consistently performing a more probable action (i.e., picking majority color balls), there would be no reason to revise the predictive model. In order to test this assumption, we analyzed the differences in pupillary responses right before and after observing a change in the agent’s picking behavior. Using a repeated measures ANOVA with condition (*Minority-first* and *Majority-first*) as a between-subjects factor and trials (4 and 5) as a within-subjects factor, we analyzed the pupillary responses of the 24-month-olds in different conditions on the last two trials. As predicted according to the third hypothesis, data revealed a significant interaction between condition and trials (*F* (1, 39) = 4.46, *p* = .04, *η*^2^ = 0.09). Follow-up t-test analyses showed that in the fourth trial, there was no significant difference in pupillary responses between the *Minority-first* condition (*M* = .16, *SD* = .20) and the *Majority-first* condition (*M* = .07, *SD* = .28), *t* (43) = 1.21, *p* = .23. However, in the fifth trial, pupil dilation in the *Minority-first* condition (*M* = .27, *SD* = .16) increased significantly, whereas this was not the case for the *Majority-first* condition (*M* = .09, *SD* = .22), *t* (38.65) = 3.01, *p* < .01. This finding is crucial, as it shows that toddlers combined observed outcomes and the information about the prior probability of an event with unobserved agent characteristics to revise their predictive models of the agent’s sampling actions over time. They thus showed increased response when this prediction was violated, which was assumed in the third hypothesis, but not in the other two hypotheses (see Figure 1C and Figure 3).

## Discussion

In this study, we investigated how 18-month-old infants and 24-month-old toddlers build predictive models of others’ actions. We defined three explanatory variables that one might use when predicting the actions of another person: (1) previously observed events, (2) the prior probability of certain events, and (3) the characteristics of an agent. These three variables can be ordered with respect to their complexity. The first one only involves simple change detection, the second one uses statistical inference and the third one requires processing of unobservable agent information. Accordingly, we developed three hypotheses each including one relevant variable more than the previous, thus building up in their level of processing complexity. We tested these hypotheses in an experiment in which young children observed a puppet picking colored balls one by one from a population box. Their pupillary responses were measured after each sampling event and were assumed to be an index of prediction errors.

Our findings showed that 24-month-old toddlers integrate the prior probability and current observations as well as an agent’s biases to build predictive models of an agent’s sampling actions, and thereby supported the third hypothesis. Because we measured the pupillary responses after each sampled outcome separately, we were able to monitor how these models and the resulting prediction errors evolved over time. Toddlers showed significantly larger pupillary responses when they observed an improbable as compared to a probable sampling action in the first two trials. This finding is in line with the assumption that young children form predictions based on the available statistical information: they predict that the distribution in a sample will reflect the distribution in the population from which they are drawn (Xu & Denison, 2009; Xu & Garcia, 2008) and are thus surprised when this prediction is violated.

Moreover, our findings suggest that repeated observations of the improbable outcome allowed toddlers to revise their predictions. They no longer assumed that the agent’s actions were random, but rather that they reflected a picking bias. As the agent consistently performed an improbable action, they expected the agent to keep showing this picking bias. However, when they observed that the last pick differed from the previous ones, this prediction was violated. The resulting prediction error caused a larger increase in their pupillary responses in the last trial in the *Minority-first* condition, as compared to the *Majority-first* condition. In the *Majority-first* condition, the overall sample resembled the distribution in the population box. Because this outcome was highly probable, the fact that the last ball had a different color did not lead to a strong increase in the pupillary response. These findings show that young children combined information about the prior probability of an event, observed outcomes and agent characteristics to form predictive models of the agent’s sampling actions.

We conclude that 24-month-olds integrate agent information into their predictive models. When a sample becomes highly improbable given the distribution in the population, they no longer assume that the sampling is random. Rather, they assume that the sampling is driven by specific characteristics of the agent: for some reason, the agent deliberately selects one color. For example, this reason could be a preference for one color over the other or a task that has been given to this agent.

Whereas the 24-month-olds in our study showed clear indications of integrating previously observed events, prior probability information and the agent characteristics, pupillary responses of 18-month-olds were inconclusive. It is likely that 18-month-old infants lack the sophisticated generative models that would allow them to perform on this task the way the older age group did. For example, in a functional near infrared spectroscopy (fNIRS) study, Emberson, Richards, and Aslin (2015) observed neural responses to violation of expectations in 6-month-olds, which suggests that the basic neural mechanisms for the generation of predictions are already in place early on in life. However, the authors also suggest that the internal models on which these predictions are based are not as sophisticated as those of adults. Similarly, this might explain the differences between 16-month-old infants and 24-month-old toddlers in reasoning about the subjective nature of preferences in the study by Ma and Xu (2011). Overall, predictive models of the environment may get more mature over the course of development allowing children to predict current input and upcoming events better.

If we apply this idea to our experiment, the 18-month-old infants’ models might not be advanced enough to integrate all three explanatory variables and the interactions between them. Alternatively, even if their internal models were advanced, they may not have generated very precise predictions resulting in low weighting on the prediction errors. For example, based on their internal model, infants may have a vague idea that what happened before is likely to happen again (cf. Hypothesis 1), but this idea might have been too weak to generate a prediction with high precision. Therefore, no or only a very weak prediction error would arise if this prediction is violated. Furthermore, it could even be the case that their internal models did not incorporate the causal link between the agent and the appearance of the balls, potentially preventing them from encoding the relevance of the color of the balls. Therefore, because of immature internal models, infants could have had weaker predictions, incorrect predictions or no predictions at all, all of which could explain the lack of overlap between their pupil response data and our hypotheses.

Predictive models of the environment might get more mature over the course of development, allowing children to make precise or detailed predictions about events. In daily life, children experience many events and most of the time there is a structure in these events. Certain events follow each other, which enables them to learn the regularities in the environment, and eventually the causal structure behind events (Gopnik, 2012). Repeated experiences of certain events might allow them to improve their model of the world to make more precise or detailed predictions. With many of these experiences, a general model of the world develops. As children gather evidence on many different occasions involving a variety of agents and objects they choose, they collect more and more world knowledge. For example, they might see a friend repeatedly picking strawberries rather than pears or their father taking coffee rather than tea. All these experiences together might allow them to integrate new information in their world model efficiently. In other words, as their world knowledge improves, they become better at inferring the causes of others’ behavior without observing many occurrences. As a result, toddlers might need less information to form a certain prediction about other agents’ choices. Potentially, 24-month-olds have gathered the world knowledge necessary to be able to use the three explanatory variables in our experiment in the way specified in the third hypothesis. These findings shed light upon the mechanisms behind toddlers’ inferences about agent-caused events. They suggest that from 24 months of age relevant information from the environment is used to form predictions about the causes of these events. Moreover, these findings show that predictive processing, which has been suggested to be a general framework explaining brain functioning, provides promising explanations for the way in which young children process another person’s actions.

## Conclusions

We investigated the computational mechanism behind young children’s inferences about an agent’s biases. We presented formalized hypotheses of how they combine perceptual, statistical and agent-related information to form predictive models of others’ actions. Our findings support the hypothesis that 24-month-old toddlers are able to integrate information about individual agents with information about previous events and prior probabilities to generate predictive models of others’ actions. Moreover, we present an innovative approach in which young children’s pupillary responses are used as indirect behavioral markers of prediction errors, as described in the predictive processing framework. The pattern of pupillary responses in 24-month-olds, but not 18-month-olds, showed strong similarities with the prediction error patterns formalized by a predictive processing model. Our findings suggest that predictive processing framework provides promising explanations of the way in which young children process another person’s actions.

## Acknowledgements

The European Union Seventh Framework Program Initial Training Network ACT (289404) and an NWO-TOP grant (407-11-040) to H.B. and I.v.R. supported this work. We would like to thank the children and families who participated in this study. We thank Gustaf Gredebäck for his advice on pupil data analyses. We would also like to thank Emil J. Wagner and Angel Cano for their help in editing the stimulus movies.

## Appendix

### Computational Model

We formalized the three hypotheses described in the introduction of this paper in a set of computational models. These computational models, based on the *causal Bayesian network* formalization of predictive processing as proposed in Kwisthout, Bekkering, and van Rooij (2017), compute posterior probability distributions that represent the expectation or *prediction* of the infant prior to each ball drawn. This prediction is compared to the actually observed event, yielding the prediction error. The *size* of this prediction error, quantified as the Kullback-Leibler divergence between the observed and predicted probability distributions (Kullback & Leibler, 1951), is a qualitative proxy for the pupil dilation. The prediction error is instrumental in updating the current beliefs that give rise to future predictions. These sub-processes in predictive processing (prediction, observation, prediction error computation, and belief updating) are graphically depicted in the context of the experimental paradigm in Figure 5.

**Figure 5:**
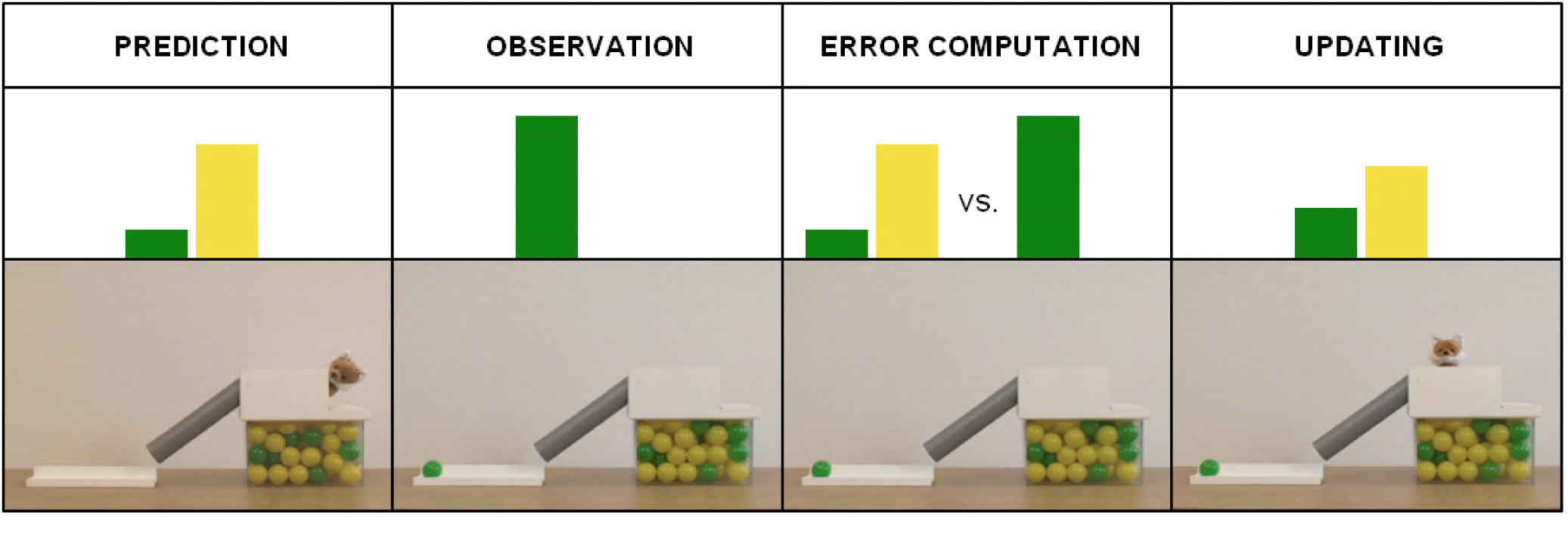
Sub-processes in predictive processing: prediction, observation, prediction error computation, and belief updating. Green and yellow bars represent probability distributions.

In the predictive processing account, the brain continuously predicts its inputs using generative models that represent the causal structure of the world (Clark, 2013). In our computational characterization, these generative models take the form of causal Bayesian networks (Pearl, 2000) with hypothesis nodes (representing the potential causes of the phenomena observed), prediction nodes (representing the observable information), and intermediate nodes (representing contextual information). The attributed *preference* of the agent is represented as a hypothesis node, the predicted *outcome* of the draw is represented as a prediction node, and the *container distribution* and the *previous draws* are represented as intermediate nodes. In the model representing Hypothesis 1, the container distribution and the preference of the agent is absent, and predictions are solely made based on the previous balls drawn. In the model representing Hypothesis 2, the container distribution is included, but the preference of the agent is still absent. In the model representing Hypothesis 3, finally, all three aspects are included. Figure 6 graphically depicts the structure of these three models prior to drawing one of the balls.

**Figure 6:**
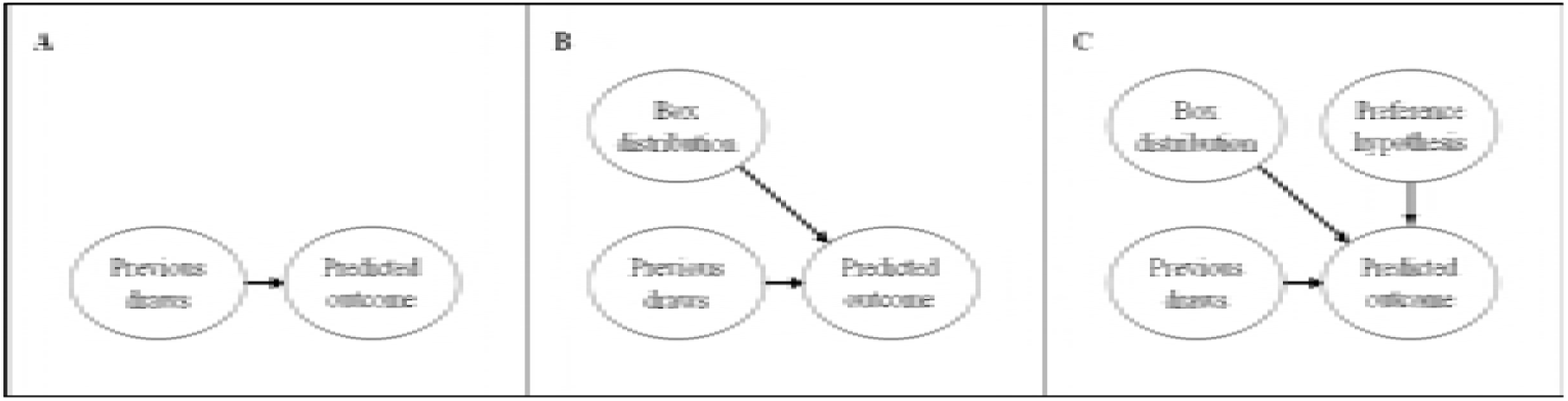
In the left panel, the model for H1 is depicted. Preceding every ball that is drawn from the box, the prediction (represented by the prediction variable Pred) is based on the previous ball (intermediate variable Prev) only; a uniform distribution is assumed for the first ball. There are no hypotheses regarding picking bias; also, the content of the container is not modeled, as it is ignored in H1. The model for H2 is depicted in the middle panel. Here, in addition to the information regarding the previous ball, the contents of the container form a contextual influence that modulates the prediction; here modeled as an additional intermediate variable, that is, Box distribution. In the right panel, the model for H3 is depicted, in which in addition to the available contextual information also the attributed agent preference is included. As this preference is a causal explanation for the ball drawn - rather than a contextual influence –, which can be updated in the light of prediction errors, we model this variable as a hypothesis variable Hyp.

With respect to the conditional probability distributions, we assume that the Box distribution variable represents the statistics of the container, that is, P (majority color) = 0.8 and P (minority color) = 0.2. We ignore the (relatively minimal) changes in the container over time and keep these prior probabilities constant. In the model corresponding with H1, we define P (Pred = C | Prev = C) = 0.8 for both the minority and majority color C; with P (Pred = minority color) = P (Pred = majority color) = 0.5 before any ball is drawn. In H2, we condition on the Box distribution variable as well and have P (Pred = C | Prev = C, Box = C) = 0.8, P (Pred = C | Prev = C, box = ¬C) = 0.2, and P (Pred = C | Box = C) = 1; here, ¬C denotes the opposite color, that is, if C is the minority color, ¬C is the majority color and vice versa. Note that in our model the previous color is always observed and that P (Pred = ¬C) = 1 – P (Pred = C), i.e., this fully defines the conditional probability distribution. In H3, this model still holds for the majority case, but we also condition on the attributed preference as well. We define P (Pred | Prev_t_ = P, Box = B, Hyp = H), where Prev_t_ [with *t* = 1…4] denotes the t-th ball drawn, as follows:

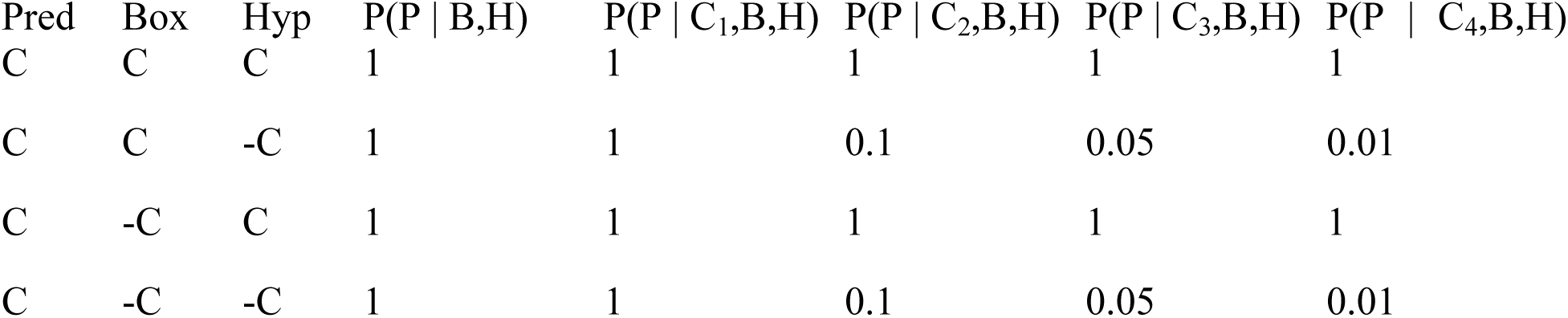

Again note that P (Pred = ¬C) = 1 - P(Pred = C) such that this table fully describes the probability distribution for the situation where C1 = C2 = C3 = C4 is the minority color. Given these computational models, we can compute the prediction error for each condition (*Minority-first/Majority-first*) and for each of the three hypotheses before every ball drawn. These prediction errors are depicted in Figure 1 in the introduction.

## References

Addyman, C., Rocha, S., & Mareschal, D. (2014). Mapping the origins of time: Scalar errors in infant time estimation. Developmental Psychology, 50(8), 2030. doi: 10.1037/a0037108.

Aslin, R. N. (2007). What’s in a look? Developmental Science, 10(1), 48–53. doi: 10.1111/j.1467-7687.2007.00563.x.

Clark, A. (2013). Whatever next? Predictive brains, situated agents, and the future of cognitive science. Behavioral and Brain Sciences, 36(03), 181–204. doi:10.1017/S0140525X12000477.

den Ouden, H. E., Daunizeau, J., Roiser, J., Friston, K. J., & Stephan, K. E. (2010). Striatal prediction error modulates cortical coupling. The Journal of Neuroscience, 30(9), 3210–3219. doi: 10.1523/JNEUR0SCI.4458-09.2010.

Egner, T., Monti, J. M., & Summerfield, C. (2010). Expectation and surprise determine neural population responses in the ventral visual stream. The Journal of Neuroscience, 30(49), 16601–16608. doi: 10.1523/JNEUR0SCI.2770-10.2010.

Emberson, L. L., Richards, J. E., & Aslin, R. N. (2015). Top-down modulation in the infant brain: Learning-induced expectations rapidly affect the sensory cortex at 6 months. Proceedings of the National Academy of Sciences, 112(31), 9585–9590. doi: 10.1073/pnas.1510343112.

Friston, K. (2005). A theory of cortical responses. Philosophical Transactions of the Royal Society B: Biological Sciences, 360(1456), 815–836. doi:10.1098/rstb.2005.1622.

Gopnik, A. (2012). Scientific thinking in young children: Theoretical advances, empirical research, and policy implications. Science, 337(6102), 1623–1627. doi: 10.1126/science.1223416.

Gopnik, A., & Wellman, H. M. (2012). Reconstructing constructivism: Causal models, Bayesian learning mechanisms, and the theory theory. Psychological Bulletin, 138(6), 1085. doi: 10.1037/a0028044.

Gredebäck, G., & Melinder, A. (2011). Teleological reasoning in 4-month-old infants: pupil dilations and contextual constraints. PloS one, 6(10), e26487. doi:10.1371/journal.pone.0026487.g001.

Henrichs, I., Elsner, C., Elsner, B., Wilkinson, N., & Gredebäck, G. (2014). Goal certainty modulates infants’ goal-directed gaze shifts. Developmental Psychology, 50(1), 100. doi: 10.1037/a0032664.

Hepach, R., & Westermann, G. (2016). Pupillometry in infancy research. Journal of Cognition and Development, 17(3), 359–377. doi: 10.1080/15248372.2015.1135801.

Jackson, I., & Sirois, S. (2009). Infant cognition: going full factorial with pupil dilation. Developmental Science, 12(4), 670–679. doi: 10.1111/j.1467-7687.2008.00805.x.

Kushnir, T., Xu, F., & Wellman, H. M. (2010). Young children use statistical sampling to infer the preferences of other people. Psychological Science, 21(8), 1134–1140. doi: 10.1177/0956797610376652.

Kullback, S., & Leibler, R. A. (1951). On information and sufficiency. The Annals of Mathematical Statistics, 22(1), 79–86.

Kwisthout, J., Bekkering, H., & van Rooij, I. (2017). To be precise, the details don’t matter: On predictive processing, precision, and level of detail of predictions. Brain and Cognition. dx.doi.org/10.1016/j.bandc.2016.02.008.

Laeng, B., Sirois, S., & Gredebäck, G. (2012). Pupillometry: a window to the preconscious?. Perspectives on Psychological Science, 7(1), 18–27. doi: 10.1177/1745691611427305.

Ma, L., & Xu, F. (2011). Young children’s use of statistical sampling evidence to infer the subjectivity of preferences. Cognition, 120(3), 403–411. doi: 10.1016/j.cognition.2011.02.003.

Meltzoff, A. N., & Gopnik, A. (2013). Learning about the mind from evidence: Children’s development of intuitive theories of perception and personality. In S. Baron-Cohen, H. Tager-Flausber, & M. Lombardo (Eds.), Understanding other minds (3rd ed., pp. 19–34). Oxford, England: Oxford University Press.

Nassar, M. R., Rumsey, K. M., Wilson, R. C., Parikh, K., Heasly, B., & Gold, J. I. (2012). Rational regulation of learning dynamics by pupil-linked arousal systems. Nature Neuroscience, 15(7), 1040–1046. doi: 10.1038/nn.3130.

Pearl, J. (2000). Causal inference without counterfactuals: Comment. Journal of the American Statistical Association, 95(450), 428–431.

Phillips, H. N., Blenkmann, A., Hughes, L. E., Bekinschtein, T. A., & Rowe, J. B. (2015). Hierarchical organization of frontotemporal networks for the prediction of stimuli across multiple dimensions. The Journal of Neuroscience, 35(25), 9255–9264. doi: 10.1523/JNEUR0SCI.5095-14.2015.

Preuschoff, K., ’t Hart, B. M., & Einhauser, W. (2011). Pupil dilation signals surprise: evidence for noradrenaline’s role in decision making. Frontiers in Neuroscience, 5, 115. doi: 10.3389/fnins.2011.00115.

Téglás, E., Girotto, V., Gonzalez, M., & Bonatti, L. L. (2007). Intuitions of probabilities shape expectations about the future at 12 months and beyond. Proceedings of the National Academy of Sciences, 104(48), 19156–19159. doi: 10.1073/pnas.0700271104.

van Pelt, S., Heil, L., Kwisthout, J., Ondobaka, S., van Rooij, I., & Bekkering, H. (2016). Beta- and gamma-band activity reflect predictive coding in the processing of causal events. Social Cognitive and Affective Neuroscience, 11(6), 973–980. doi: 10.1093/scan/nsw017.

Wacongne, C., Labyt, E., van Wassenhove, V., Bekinschtein, T., Naccache, L., & Dehaene, S. (2011). Evidence for a hierarchy of predictions and prediction errors in human cortex. Proceedings of the National Academy of Sciences, 108(51), 20754–20759. doi: 10.1073/pnas.1117807108.

Wellman, H. M., Kushnir, T., Xu, F., & Brink, K. A. (2016). Infants use statistical sampling to understand the psychological world. Infancy, 21(5), 668–676. doi: 10.1111/infa.12131.

Xu, F., & Denison, S. (2009). Statistical inference and sensitivity to sampling in 11-month-old infants. Cognition, 112(1), 97–104. doi:10.1016/j.cognition.2009.04.006.

Xu, F., & Garcia, V. (2008). Intuitive statistics by 8-month-old infants. Proceedings of the National Academy of Sciences, 105(13), 5012–5015. doi: 10.1073/pnas.0704450105.

